# BuTT-Seq: a new method for facile profiling of transcription

**DOI:** 10.1101/2023.03.04.531084

**Authors:** Albert D. Yu, Michael Rosbash

## Abstract

A wide range of sequencing methods have been developed to assess nascent RNA transcription and resolve the single-nucleotide position of RNA polymerase genome-wide. These techniques are often burdened with high input material requirements and lengthy protocols. We leveraged the template-switching properties of thermostable group II intron reverse transcriptase (TGIRT) and developed BuTT-Seq (BUlk analysis of nascent Transcript Termini sequencing), which can produce libraries from purified nascent RNA in 6 hours and from as few as 10,000 cells – an improvement of at least 25-fold over existing techniques. BuTT-Seq shows that inhibition of the superelongation complex (SEC) causes promoter-proximal pausing to move upstream in a fashion correlated with subnucleosomal fragments. To address transcriptional regulation in a tissue, BuTT-Seq was used to measure the circadian regulation of transcription from fly heads. All the results indicate that BuTT-Seq is a simple and powerful technique to analyze transcription at a high level of resolution.

## INTRODUCTION

Although the analysis of messenger RNA (mRNA) can usually be done with simple RNA-Seq methods, the analysis of transcription and nascent RNA is faced with greater challenges. Not only is nascent RNA less abundant, but it is also substantially more heterogeneous than mRNA; nascent RNA contains diverse transcripts beyond pre-mRNA, and these transcripts are in various states of active or stalled synthesis. This complexity requires careful consideration.

There are three general classes of techniques that analyze transcription. The first, Chromatin Immunoprecipitation and Sequencing (ChIP-Seq), immunoprecipitates RNA Polymerase II (RNAPII) and sequences the associated DNA fragments. Such a standard ChIP-Seq approach provides a low resolution view of RNAPII localization, although single-nucleotide positioning can be achieved with modifications such as ChIP-Nexus (He et al., 2015). The second class, run-on techniques such as global run-on sequencing (GRO-Seq) and precision run-on and sequencing (PRO-Seq), isolate nuclei containing RNA undergoing active synthesis. Transcription is then begun in vitro with tagged nucleotides that can be affinity purified and sequenced(Core et al., 2008; Mahat et al., 2016). Whereas these run-on techniques feature the greatest selectivity for actively synthesized RNA, they cannot capture RNA in stalled synthesis complexes, such as those with backtracked RNAPII (Wissink et al., 2019). The third class of techniques relies on the biochemical purification of nascent RNA and therefore captures RNA in both active and stalled RNAPII complexes.

The remarkable stability of the RNA Polymerase/nascent RNA/DNA ternary complex enables the isolation of nascent RNA with buffers containing a high concentration of salt and urea (Wuarin & Schibler, 1994). Techniques leveraging biochemically purified nascent RNA include preparing libraries from the entire length of the nascent RNA molecule, such as with Nascent-Seq or chromatin-associated RNA sequencing (chrRNA-seq) (Khodor et al., 2012; Sousa-Luis et al., 2021). Other techniques aim to track RNA synthesis at single-nucleotide resolution by ligating adapters to and sequencing the 3’ ends of nascent RNA molecules; the logic is that the 3’ end is the most recently synthesized base. These 3’end techniques include short capped RNA sequencing (scRNA-seq), 3’ nucleotide sequencing (3NT-seq), and human native elongating transcript sequencing (hNET-Seq) (Mayer et al., 2015; Nechaev et al., 2010; Weber et al., 2014). scRNA-seq and 3NT-seq also feature the enrichment of capped RNA by digestion of uncapped RNA with terminator exonuclease. A variation of 3’ end sequencing techniques involves the immunoprecipitation of RNAPII to enable phosphorylation-state specific profiling of nascent RNA. They include native elongating transcript sequencing (NET-Seq) and its derivatives, e.g., mammalian NET-Seq (mNET-Seq) and enhanced NET-Seq (eNET-Seq) (Churchman & Weissman, 2012; Fong et al., 2022; Nojima et al., 2015).

These powerful techniques that sequence biochemically purified nascent RNA also confront some contamination issues, for example by mature chromatin-associated RNAs like snoRNAs. Thus, care must be taken when interpreting signal. Moreover, all of these methods share two less than ideal features: the protocols are relatively complex and time-consuming, and they have quite high input material requirements.

To circumvent these features, we have devised a novel approach towards sequencing the 3’ends of biochemically purified nascent RNA, which we have named BuTT-Seq (**<u>Bu</u>**lk analysis of nascent **<u>T</u>**ranscript **<u>T</u>**ermini sequencing). BuTT-Seq leverages an optimized Thermostable Group II Intron Reverse Transcriptase (TGIRT) reverse transcription protocol, which couples reverse transcription and adapter ligation into a single step (Mohr et al., 2013). This protocol is used for both first and second synthesis with several optimizations, such as a salt-concentration switching step to minimize template switches and maximize RT efficiency and a low concentration of dNTPs to minimize fragment size. These steps eliminate the need for RNA fragmentation as well as multiple materially demanding and time-consuming gel extraction steps. BuTT-Seq can prepare libraries from purified nascent RNA in as few as 6 hours and exhibits reproducible signal with inputs as low as 10,000 *Drosophila* S2 cells. This input requirement represents at least a 25-fold improvement over existing protocols; other NET-Seq like techniques typically reference inputs of 1,000,000-10,000,000 cells, while qPRO-Seq requires 250,000 cells (Churchman & Weissman, 2012; Judd et al., 2020; Mayer et al., 2015; Nojima et al., 2015).

BuTT-Seq can resolve RNA polymerase localization at single nucleotide resolution: pharmaceutically inhibiting the SEC with KL-1 causes pause sites to recede upstream, implying that promoter-proximal pausing regulation may involve the clearance of multiple discrete pausing sites. BuTT-Seq also is effective with limited input material, which we demonstrate by profiling small numbers of *Drosophila* heads. Altogether, we establish BuTT-Seq as a simple and powerful method capable of resolving multiple aspects of transcription that offers several attractive improvements over existing methods.

## RESULTS

### BuTT-Seq is an efficient 3’ Nascent RNA end-labeling technique

Nascent RNA contains a heterogeneous mixture of molecules in varying states of synthesis. They include short transcripts associated with paused RNA polymerase, partially synthesized RNAs, and completed transcripts awaiting cleavage and polyadenylation (Wissink et al., 2019). BuTT-Seq is inspired by other 3’-end labeling techniques and captures the entire breadth of all nascent transcripts. However, the data presented focus on transcripts from protein-coding genes.

The protocol first isolates chromatin with 1M Urea and 3% Empigen as previously described (Rodriguez et al., 2013; Schlackow et al., 2017; Sousa-Luis et al., 2021). The resulting chromatin-associated RNA undergoes simultaneous adapter ligation and reverse transcription using the TGIRT reverse transcriptase and an adapter containing an 8-bp UMI in place of the usual i7 index. A short reverse transcription step is conducted with a low concentration of dNTPs; this is to limit the length of the cDNA and minimize length-dependent biases when sequencing (Figure 1A). Furthermore, pre-RT incubation is done with a low concentration of salt, while dNTPs are added along with a higher concentration of salt; this is done to maximize the efficiency of the initial template switch, but to minimize the amount of secondary template switches as well as slow down reverse transcription. Second-strand synthesis and adaptor ligation are conducted through a second TGIRT reaction, and the final adapter-tagged cDNA is subject to PCR with standard i5 indexing primer, gel extraction, and standard Illumina sequencing. The resulting library should reflect all chromatin-associated RNA, most notably a high-resolution view of polymerase pausing and transcriptional elongation (Figure 1A).

**Figure 1.**
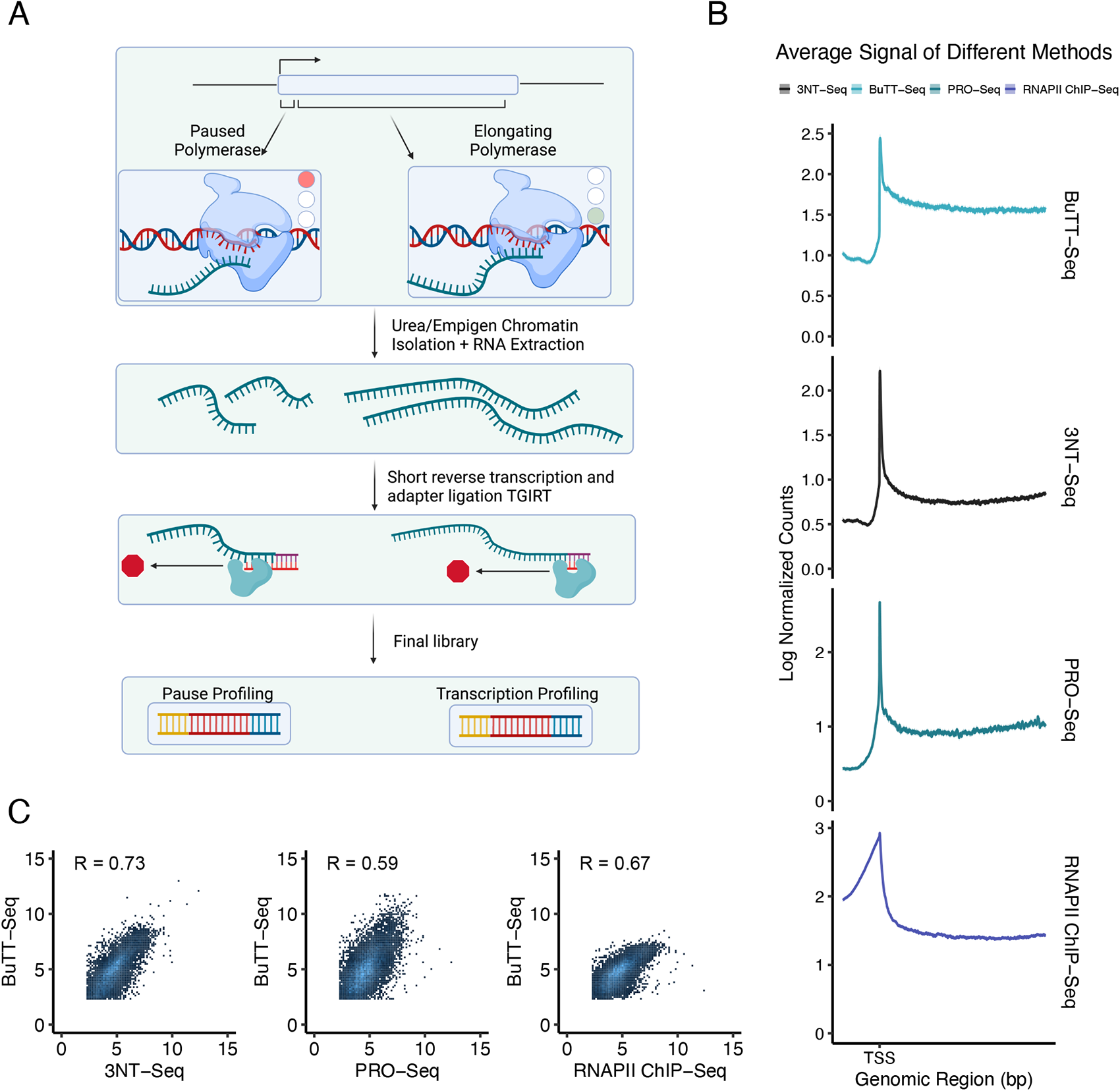
BuTT-Seq measures transcription and is comparable to other transcription analysis methods. (A) BuTT-Seq method overview. (B) Scaled metagene plots of signal distribution in plus strand genes over 5kb in length (N=6284) across BuTT-Seq, 3NT-Seq, PRO-Seq, and RNAPII ChIP-Seq. On the X axis, the transcription start site (TSS) is marked and plot extends 1kb into the gene body and 200bp upstream. The Y axis represents the log2-transformed normalized signal in each method. The shaded area corresponds to the 95% confidence interval. (C) Log-log plot of normalized counts from BuTT-Seq compared to 3NT-Seq, PRO-Seq, and RNAPII ChIP-Seq genome-wide(N=. Signal was quantified from the region 200nt downstream of the TSS genome-wide. The Y-axis represents the log2-transformed BuTT-Seq counts, while the X-axis represents the log2-transformed 3NT-Seq(Left), PRO-Seq(Middle), and RNAPII ChIP-Seq(Right) counts. Correlations were calculated using Spearman’s rank correlation coefficient.

### BuTT-Seq results are comparable to those from other established methods

A major use of 3’-end labeling techniques is the identification and characterization of promoter-proximal pausing, represented by regions of distinctly high signal adjacent to the transcription start site (Mayer et al., 2017). Such pausing represents a distinct regulatory step: RNAPII transcribes a short stretch of RNA before pausing and awaiting further signaling before either disengaging or proceeding into processive elongation. To validate BuTT-Seq for pausing analysis, we examined the average signal distribution 200nt downstream of the TSS genome-wide in *Drosophila* S2 cells and compared it to the signal from published data in these cells generated by other nascent RNA techniques: 3NT-Seq, PRO-Seq, and RNAPII ChIP-Seq (Figure 1B). All four techniques feature a sharp peak directly downstream of the TSS, likely driven by the high prevalence of promoter-proximal pausing across the genome, followed by signal reflecting transcripts undergoing processive elongation.

We quantified these regions and performed a log-log comparison of BuTT-Seq against the three published techniques. BuTT-Seq has the highest correlation to the most chemically analogous technique, 3NT-Seq (Spearman correlation = 0.73), but also correlates well with RNAPII ChIP-Seq (Spearman’s correlation = 0.67) and PRO-Seq (Spearman’s correlation = 0.59). These comparisons indicate that BuTT-Seq faithfully assays nascent RNA.

### BuTT-Seq is reproducible down to 10,000 cells

We next assayed the minimum input requirement for BuTT-Seq. Our eventual goal is to assay primary tissues or cells from which material may be limited. The BuTT-Seq signal 200nt downstream of the TSS correlated well between 500,000 and 50,000 cells (Spearman’s correlation = 0.97), and even between 500,000 and 10,000 cells (Spearman’s correlation = 0.91) (Figure 2A). There is a similar signal distribution pattern: all feature a sharp peak at the TSS indicative of promoter-proximal pausing (Figure 2B). The high correlation coefficients indicates that BuTT-Seq is not only highly reproducible but also consistent with 10,000 cells or less.

**Figure 2.**
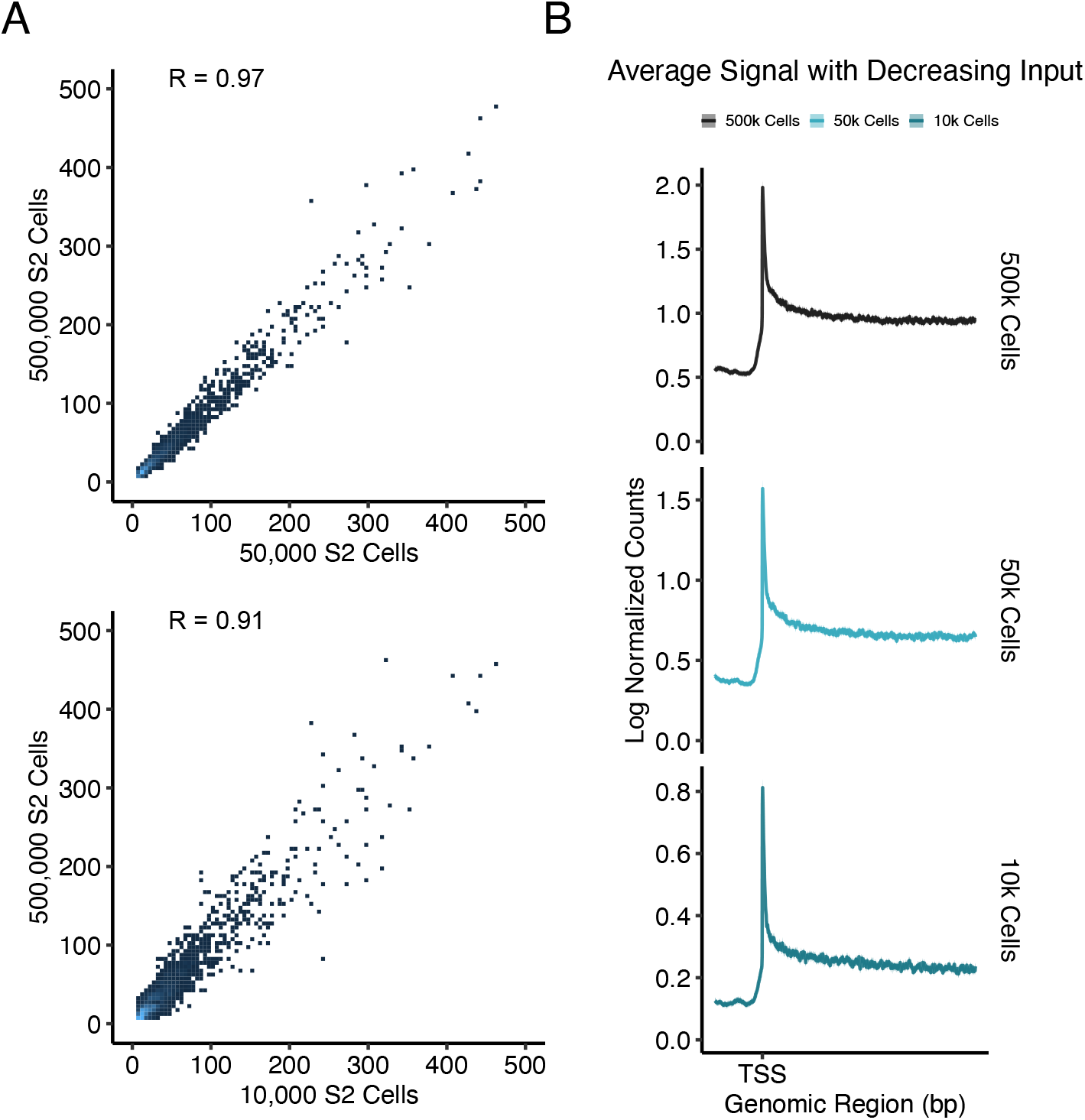
BuTT-Seq signal is reproducible down to 10,000 cells. (A) Comparison of signal from BuTT-Seq data generated from 500,000, 50,000, and 10,000 cells. Signal was quantified from 200nt downstream of the TSS genome-wide. The Y-axis represents the log2-transformed counts from 500,000 cells, while the X-axis represents the log2-transformed counts from 50,000 (Top) and 10,000 (Bottom) cells. (B) Metagene plots of signal distribution in plus strand genes over 5kb in length (N=6284) from 500,000, 50,000, and 10,000 cells. On the X axis, the transcription start site (TSS) is marked and plot extends 1kb into the gene body and 200bp upstream. The Y axis represents the log2-transformed normalized signal in each method. The shaded area corresponds to the 95% confidence interval.

### KL-1 causes pause sites to recede towards the TSS

We next used BuTT-Seq to characterize the effect of the AFF4/lilliputian (*lilli*) inhibitor, KL-1, on transcription in S2 cells. KL-1 disrupts the formation of the superelongation complex (SEC) by inhibiting *lilli* activity (Liang et al., 2018). It thereby impedes the release of paused polymerase into productive elongation and attenuates the rate of elongation. As KL-1 has previously been shown to result in an increased RNAPII ChIP-Seq signal at the TSS, we compared BuTT-Seq signal to published RNAPII ChIP-Seq data (Liang et al., 2018).

To assay the genome-wide pausing index of 20µM or 60µM KL-1-treated S2 cells compared to vehicle (DMSO)-treated cells, we calculated the signal ratio between promoter-adjacent reads and gene body reads (Figure 3A). By measuring pausing index on an Empirical Cumulative Distribution Function (ECDF) plot, there was a significant increase in pausing index upon KL-1 treatment that was not significantly different between 20µM or 60µM, similar to what had been previously shown in RNAPII ChIP-Seq (Figure 3A) (Liang et al., 2018).

**Figure 3.**
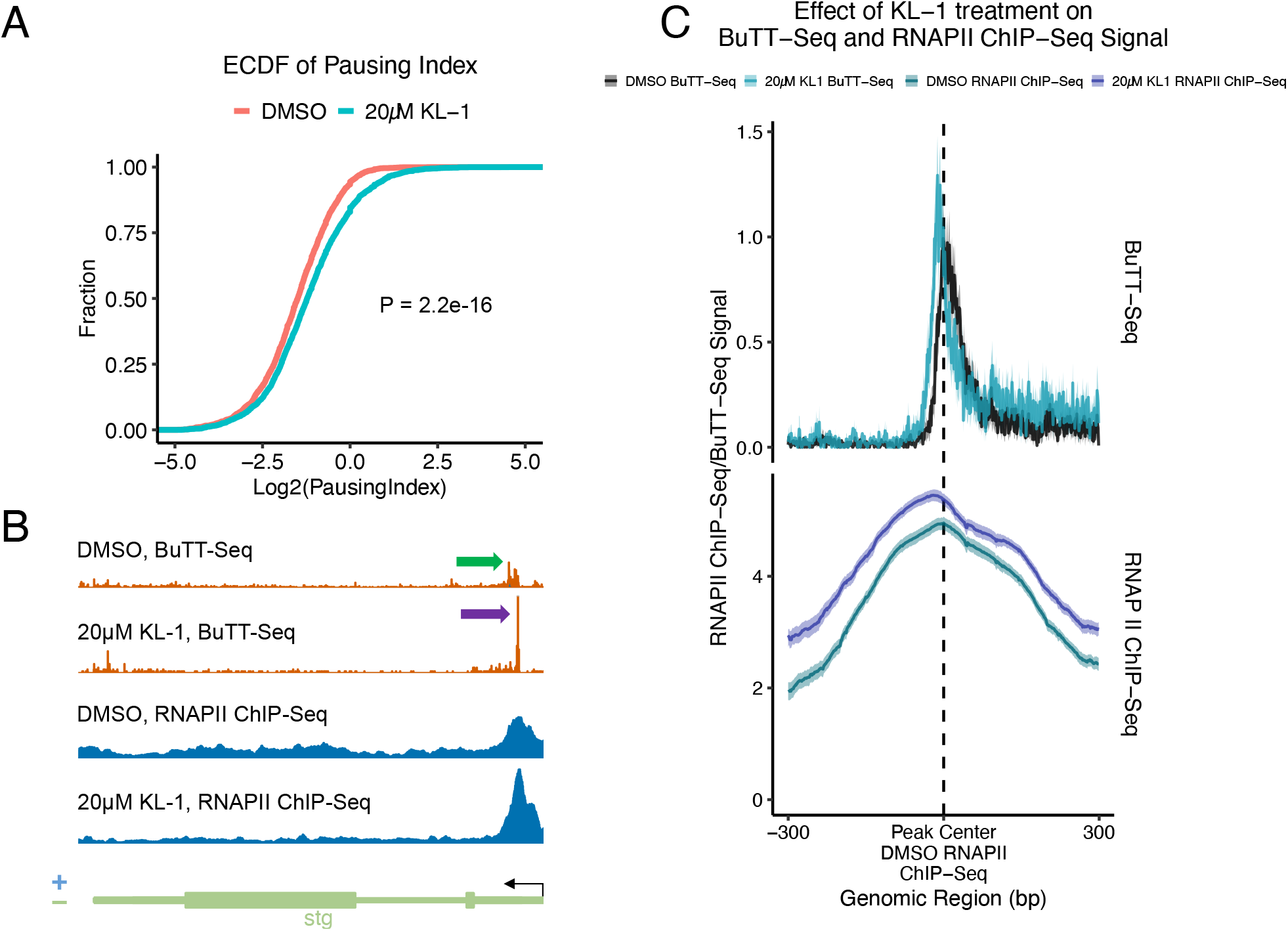
BuTT-Seq recapitulates pausing dynamics seen in RNAPII ChIP-Seq upon KL-1 treatment. (A) ECDF of pausing index in BuTT-Seq in S2 cells treated with DMSO, 20µM KL-1, or 60µM KL-1. Pausing region is defined as the region from the TSS to computationally determined pause sites, and gene body region is defined as 1000nt downstream of each pause site. The Y-axis depicts the cumulative fraction of genes, while the X-axis contains the log2-transformed pausing index. (B) Representative gene demonstrating the effect of 20µM KL-1 treatment on signal distribution in BuTT-Seq and RNAPII ChIP-Seq. Green arrow: Untreated pause site. Purple arrow: KL-1 induced pause site. (C) Metagene plots of log2-normalized BuTT-Seq and RNAPII ChIP-Seq counts, centered around RNAPII ChIP-Seq peaks called from DMSO-treated S2 cells that overlap with BuTT-Seq single-nucleotide pause peaks. Only plus strand genes are depicted (N=219). Shade region represents the 95% confidence interval.

To analyze transcriptional dynamics at single nucleotide resolution, each BuTT-Seq read was truncated to the first base sequenced by Read 2; this should reflect the last nascent RNA base synthesized by RNAPII. Surprisingly, this single-nucleotide resolution indicated that KL-1 not only increased the magnitude of pausing as previously reported but also caused genome-wide pause sites to move upstream toward the TSS (Liang et al., 2018). (A representative gene is shown in Figure 3B.) Reanalysis of published RNAPII ChIP-Seq data and metagene profiling of this RNAPII ChIP-Seq signal indicate that both of these conclusions are visible in these previous data (Figure 3C). First, the BuTT-Seq signal peaks align with the RNAPII ChIP-Seq peaks, both for DMSO and KL-1 treated conditions. Second, treatment with KL-1 causes the signal to move in the 5’ direction towards the TSS while also increasing in magnitude, suggestive of increased pausing in addition to altering the precise pause sites (Figure 3C). This effect is most pronounced in BuTT-Seq, although it is also visible at lower resolution in the RNAPII ChIP-Seq data.

### RNAPII positioning and KL-1 sensitivity correlate with subnucleosomal structures of the – 1 nucleosome

We leveraged the single nucleotide sensitivity of BuTT-Seq to address the relationship between RNAPII pausing and subnucleosome structure. MNase-seq is used to measure nucleosome positioning, and selective sequencing and analysis of sub-nucleosomal (<147bp) MNase-digested fragments can be used to map partially unwrapped subnucleosomal particles (Ramachandran et al., 2017). These fragments typically exhibit a bimodal distribution, representing protection from digestion by the two halves of a partially unwrapped histone dyad (Visible in Figure 4A, top). RNAPII proximity to the +1 nucleosome correlates with loss of the dyad distal to the TSS(Ramachandran et al., 2017). This observation was inferred to be a causal result of RNAPII transcribing through the nucleosome. However, many studies suggest the opposite: even the subnucleosome particle is a transcriptional barrier capable of slowing and pausing RNAPII, which then can be overcome by RNAPII through a number of molecular strategies. In any case, we asked whether subnucleosomal dynamics correlate with KL-1 sensitivity.

**Figure 4.**
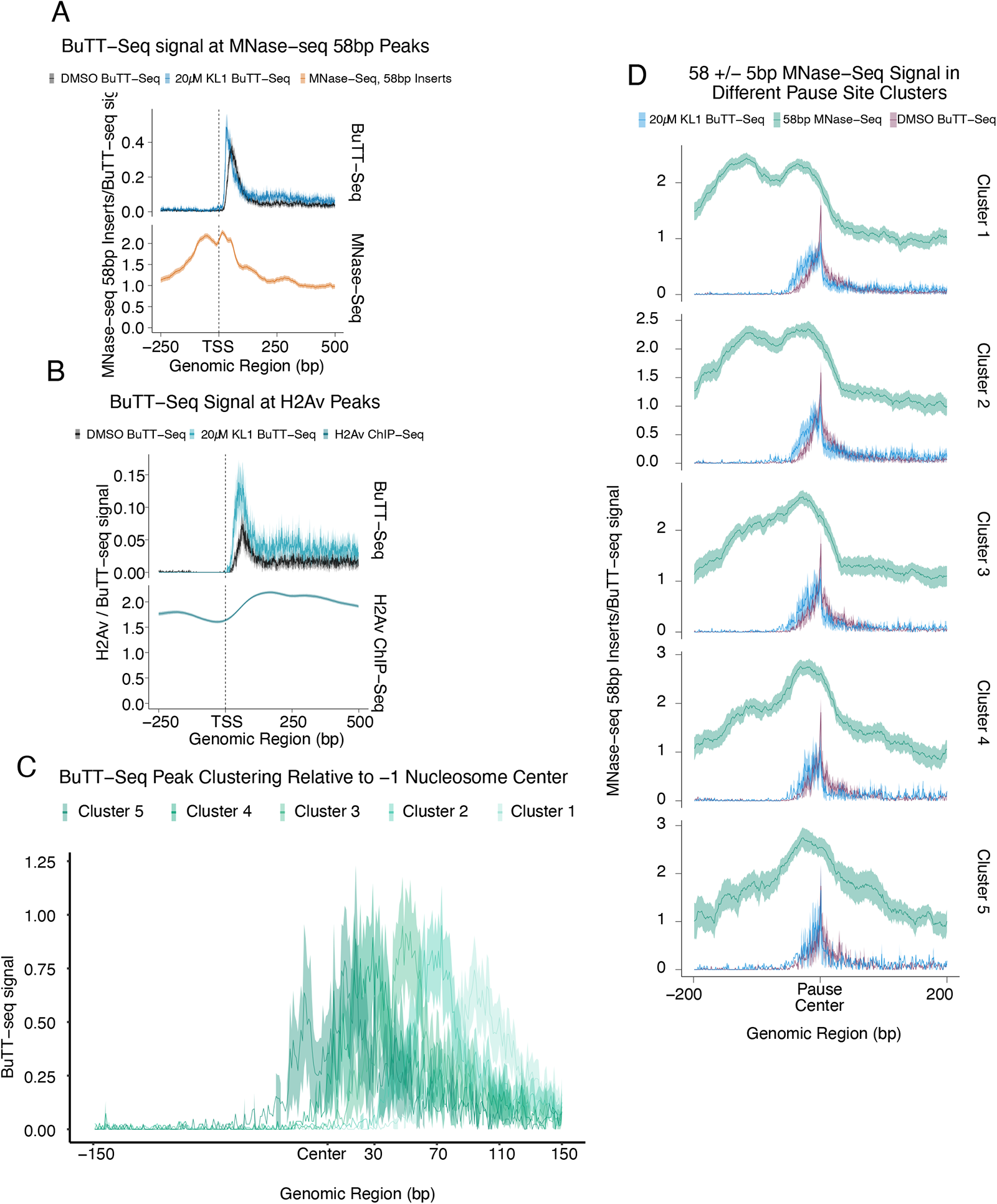
Pausing as assayed by BuTT-Seq is correlated with nucleosomal dynamics. (A) Metagene plots of log2-transformed BuTT-Seq counts compared to log2-normalized MNase-seq counts, restricted to 58bp +/-5nt fragments from MNase-seq. Plotted genes were derived from MNase-seq peaks that overlap with annotated TSS; plus-strand genes are shown (N=831). Shaded area corresponds to 95% confidence interval. (B) Metagene plot of log2-transformed H2Av ChIP-Seq counts compared to log2-transformed BuTT-Seq counts from S2 cells treated with DMSO or 20µM KL-1. Plotted genes were derived from H2Av ChIP-seq peaks that overlap with annotated TSS; plus-strand genes are shown (N=1531). Shaded area corresponds to 95% confidence interval. Clustering of pause peaks based on distance from the -1 nucleosome center. Depicted are plus strand genes. Cluster 1 contains pauses 80-120bp downstream (N=140), Cluster 2 contains pauses 60-79bp downstream (N=95), Cluster 3 40-59bp (N=63), Cluster 4 20-39bp (N=45), and Cluster 5 0-19bp. (N=35). The shaded area corresponds to the 95% confidence interval. Metagene plot of log2-normalized 58bp +/-5bp MNase-Seq counts against log2-normalized KL-1 and DMSO treated BuTT-Seq counts across 5 clusters. Shaded area corresponds to 95% confidence interval.

We first measured the genome-wide distribution of the BuTT-Seq signal and the MNase-Seq signal restricted to 58 +/-5 bp fragments (Figure 4A). A surprising number of MNase peaks with a bimodal distribution overlapped the TSS (N=1684) (visible in Figure 4A, top). The region overlapping the TSS is typically thought to be a nucleosome free-region, and the -1 nucleosome is defined as the region just preceding the TSS. However, this 58bp pattern is consistent with protection by a subnucleosomal H3-H4 dyad, a distribution referred to as a -1 nucleosome (Ramachandran et al., 2017); we will use this terminology. Pausing was enriched in genes exhibiting this fragment distribution pattern and was also sensitive to KL-1 treatment (Figure 4A, bottom). In contrast, pausing was depleted in genes enriched with the H2Av variant in the +1 nucleosome as previously shown (Figure 4B)(Weber et al., 2014). Surprisingly however, H2Av-enriched genes were also sensitive to KL-1 treatment.

To further analyze these BuTT-Seq pause sites, they were divided into 5 clusters representing their distance downstream from the center of the -1 nucleosome (Figure 4C). Cluster 1 contains pauses 80-120bp downstream of the -1 nucleosome, Cluster 2 contains pauses 60-79bp downstream, Cluster 3 contains pauses 40-59bp downstream, Cluster 4 contains pauses 20-39 downstream, while Cluster 5 contains pauses 0-19 bases downstream. Plotting MNase-Seq and KL-1 treated BuTT-Seq signals against the DMSO-treated signal in each cluster revealed two notable patterns (Figure 4D). First, the character of the nucleosome dyad changes in each cluster. In Cluster 1, where the pause site is furthest from the -1 nucleosome, both halves of the nucleosomal dyad are present. In Cluster 5, where the pause site is very close to the +1 nucleosome, the distal half of the nucleosome dyad is completely absent. Moving from Cluster 1 to Cluster 5, there is a gradual loss of the nucleosome dyad distal half (Figure 4D, blue). Second, the effect of KL-1 treatment on the pause site appears to be impacted by the distance of the pause site to the nucleosome center; the further downstream the pause site, the greater the effect of KL-This relationship

### BuTT-Seq identifies differential pausing dynamics between S2 cells and *Drosophila* heads

One of the primary advantages of BuTT-Seq is that its more relaxed input material requirements enable a more facile profiling of primary tissues. To this end, BuTT-Seq was used to profile nascent RNA from a small number (n = 5) Canton-S *Drosophila* heads and compared to the S2 cell profiles. These two very different tissues are not surprisingly poorly correlated (Spearman’s correlation = 0.30) (Figure 5A). However, many commonly expressed genes have similar single-nucleotide pausing signatures (Figure 5B).

**Figure 5.**
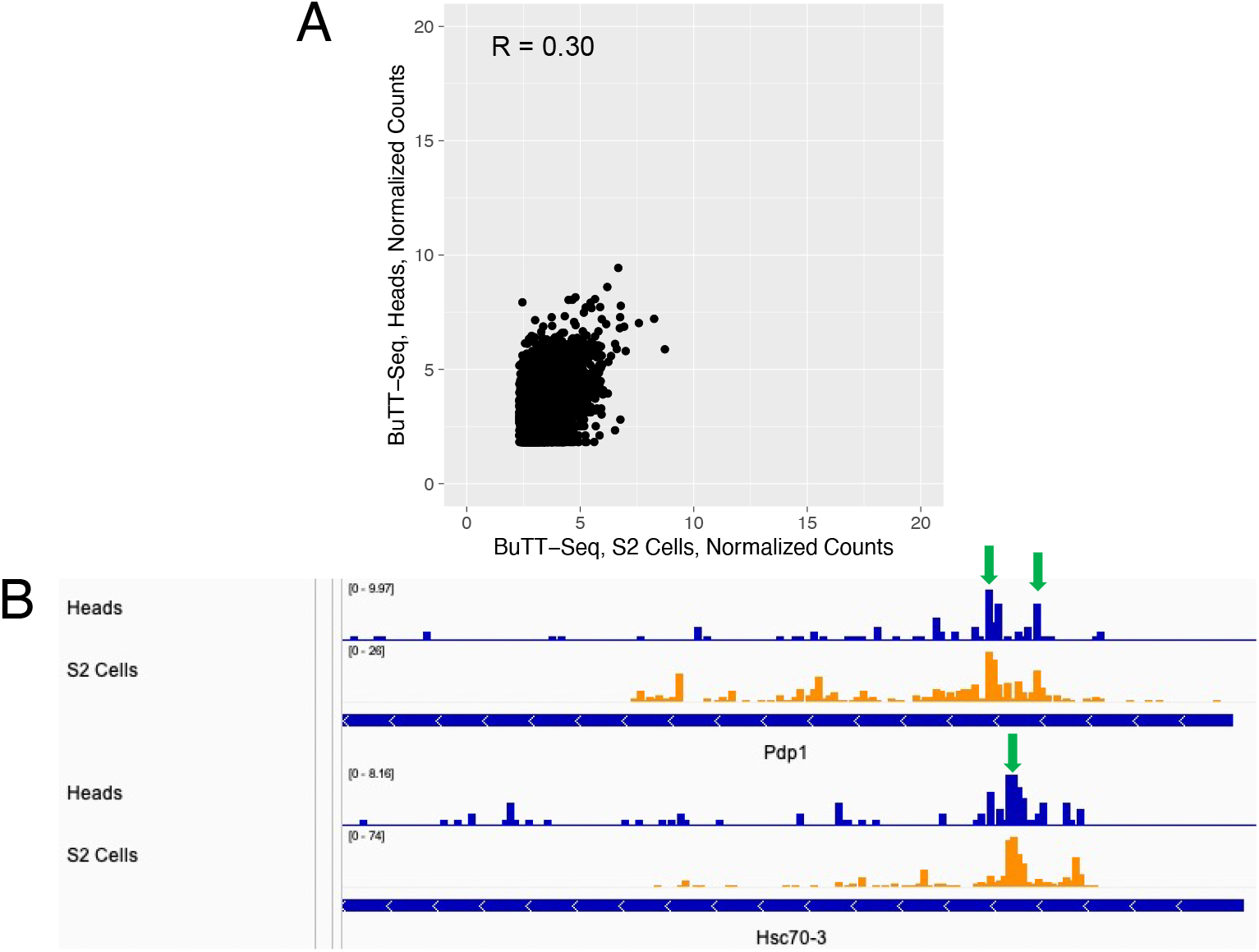
BuTT-Seq in heads bears some resemblance to S2 Cells. (A) Comparison of counts from BuTT-Seq data generated from S2 cells or heads. Signal was quantified from 200nt downstream of the TSS genome-wide. The Y-axis represents the log2-transformed counts from heads, while the X-axis represents the log2-transformed counts from S2 cells. (B) Representative genome browser view of genes exhibiting similar pause sites in Heads and S2 cells. Green arrow: Pause sites

To further explore the variation in pausing and elongation dynamics between heads and S2 cells, we first called single-nucleotide peaks separately from S2 cells and heads, restricting the peak calling to within 200nt of the TSS and filtering for the highest signal when a gene had multiple peaks. We then merged peaks between S2 cells and heads and compared signals in pausing regions and elongation regions of each gene: We defined the pause region as the region between the pause peak and the TSS and the elongation region as a gene body region 1000bp downstream of the pause peak. We then compared the relative signal between the pause region and the elongation region between S2 cells and heads (Figure 6).

**Figure 6.**
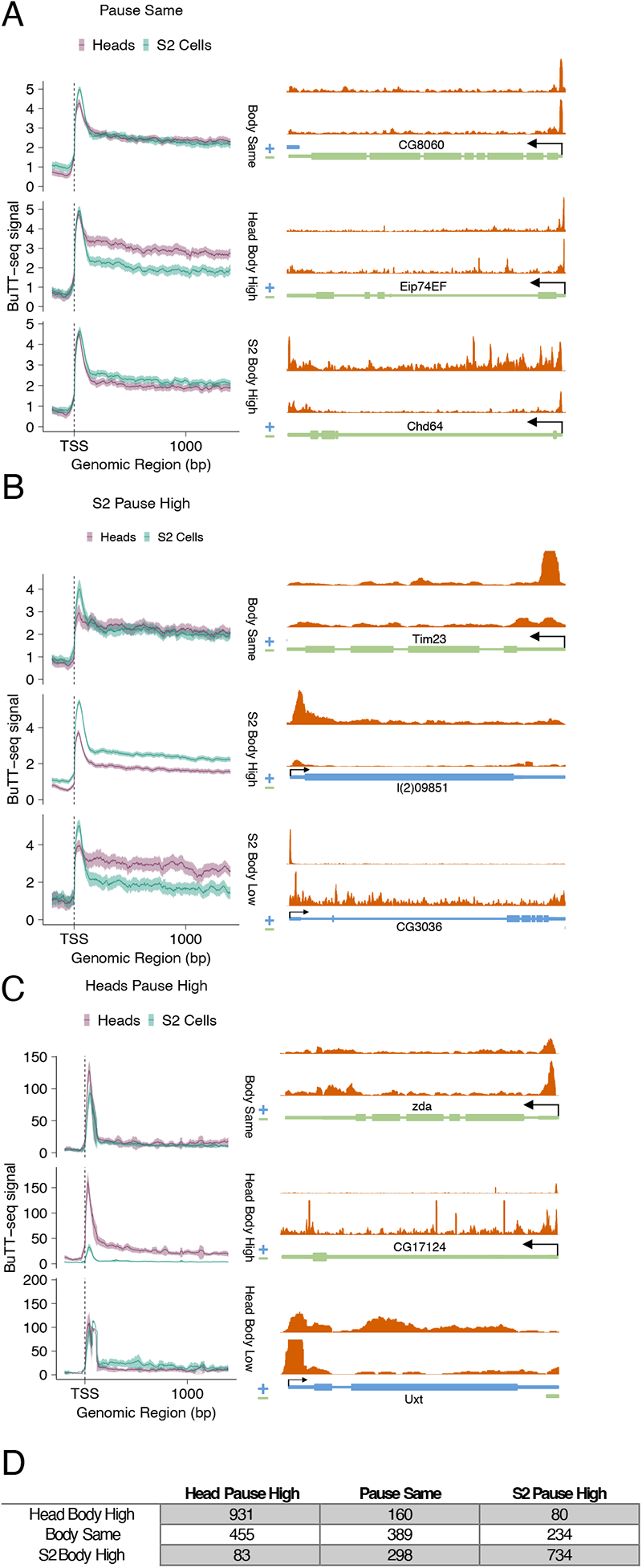
Differential analysis of BuTT-Seq in heads and S2 cells reveals a diverse range of transcriptional programs. Metagene analysis (left) and representative genes (right) reflecting different combinations of pause region and gene body signal between S2 cells and heads. (A) Pausing is the same between heads and S2 cells. (B) Pausing is higher in S2 cells. (C) Pausing is higher in heads. (D) Total number of genes occupying each pause/gene body combination.

Pausing and elongation are concordant in 61% of genes; when pausing is higher in one tissue, elongation is also higher, and vice versa (Figure 6A top, Figure 6B and 6C middle, 6D). These genes might simply manifest tissue differences in rates of transcriptional initiation. In most of the other genes (31%), there was a discordant relationship between pause and elongation regions: heads and S2 cells exhibit the same signal in one parameter, but a different signal in the other (Figure 6B and 6C, top and bottom). Most notably, 8% of genes are discordant for both parameters, e.g., when pausing is higher in one tissue, elongation is lower. These genes may exhibit differential bottlenecking by pause release factors.

### BuTT-Seq identifies transcriptional dynamics in key circadian genes

We and others have previously measured the association of RNAPII with circadian clock genes in fly heads at different times of day (Abruzzi et al., 2011), for example by ChIP-Seq. Taken together with other assays (Taylor & Hardin, 2008), there is strong evidence that RNAPII cycles on and off clock gene chromatin in gene body regions in a circadian manner and parallels cycling clock gene transcription. There is however one striking exception: there are substantial levels of RNAPII stably associated with the *period (per)* gene promoter region at all times of day, even at times of day when there is little transcription (Abruzzi et al., 2011; Taylor & Hardin, 2008). Will BuTT-Seq recapitulate or perhaps even extend these previous observations? To answer this question, we assayed fly head nuclei from entrained flies collected at 6 times of day, ZT2-ZT22 (ZT is time in a 12:12 LD cycle, with ZT0=lights on and ZT12=lights off).

As anticipated, the clock gene body signal undergoes robust circadian cycling with peaks at ZT14 and troughs at ZT2 for the CLK/CYC direct target genes *per, vri* and *tim,* and a peak at ZT2 and a trough at ZT14 for the clock gene *Clk.* (Fig. 7; only these two time points are shown). Notably, the *per* BuTT-Seq data show a prominent and stable signal across peak and trough timepoints just downstream of the transcription start site (Figure 7A top left). This indicates that pause release might be under circadian regulation and contribute to *per* transcription. In contrast, the other two direct target genes *vri* and *tim* differ from *per.* Although *tim* and *vri* show notable pause sites near their TSSs (Figure 7A top right and bottom left), they oscillate with a similar amplitude to the gene body (Figure 7A and data not shown). This suggests that the transcription of these 2 genes is primarily regulated through oscillating transcription initiation. The *Clk* gene in contrast features a much smaller but temporally constant pause site like that of *per*, which implies that *Clk* may also be at least partially regulated through pause release.

**Figure 7.**
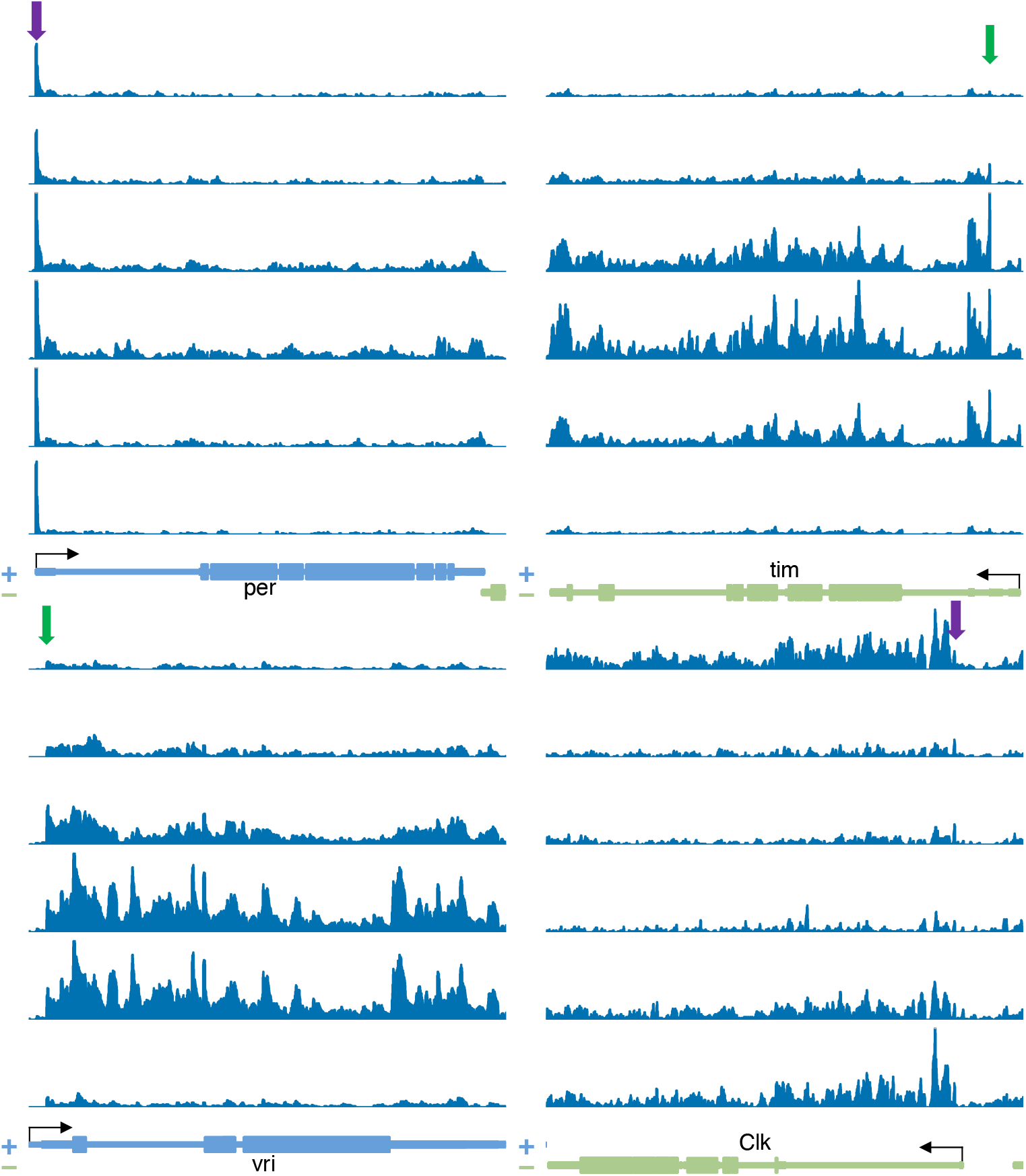
BuTT-Seq exhibits transcriptional cycling of core circadian genes. Genome browser view of six timepoints of core circadian genes in BuTT-Seq. Green Arrow: Cycling pause. Purple arrow: Constant pause.

**Figure 8.**
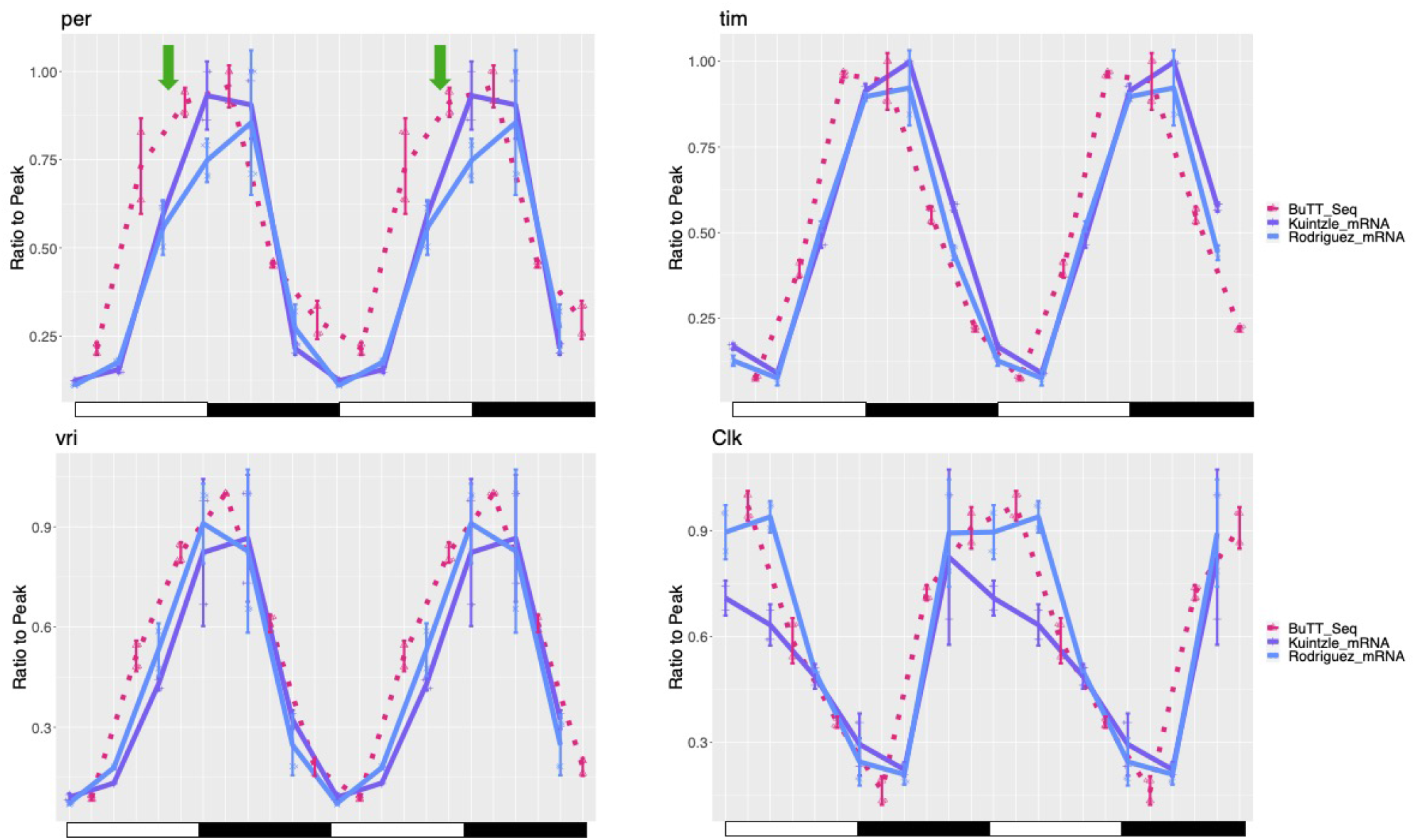
BuTT-Seq recapitulates known transcriptional features of the Circadian clock. BuTT-Seq double-plotted against RNA-Seq data from Rodriguez et al. (2013) and Kuintzle et al. (2017) at core circadian genes. Each timepoint was normalized to the peak timepoint in respective genes. Green arrow: ”Hump” of transcription in per gene.

### BuTT-Seq recapitulates circadian transcriptional oscillation details

Could BuTT-Seq be used to assess different transcription rates? Although a high BuTT-Seq signal in gene body regions may reflect a high level of RNA production, it may also reflect a high level of RNAPII occupancy due to a slow rate of transcriptional elongation. However, both NET-seq and RNAPII ChIP-Seq signal have previously been used to approximate transcription levels of RNA synthesis, which is because RNAPII occupancy often if not usually reflects RNA synthesis. To examine in quantitative detail this putative circadian transcriptional regulation, we calculated the BuTT-Seq signal within every gene at each of the six time points; only exon signals were summed, i.e., intron signals were removed to avoid possible complications from nascent splicing. To help interpret the data, we turned to two published head time point mRNA-seq datasets – one generated by our lab almost 10 years ago and the other generated elsewhere five years ago (Kuintzle et al., 2017; Rodriguez et al., 2013). For ease of illustration, we focus here on the four core clock direct CLK-target genes presented above (Figure 7A) and double plotted these two data sets along with our BuTT-Seq data (Figure 7B).

A first notable observation is that the two RNA-seq data sets are very similar; the *tim* and *vri* curves are indistinguishable, whereas only the peak values of the *Clk* profiles and to a lesser extent the *per* profiles are somewhat different (Figure 7B, purple and blue). A second is that the BuTT-Seq data are virtually superimposable onto the RNA-seq data for 3 of the four genes. This suggests that transcriptional regulation accounts for almost all mRNA dynamics for these four core clock genes (Figure 7B).

The only notable exception is *per*. The BuTT-Seq data indicate that the mid-day increase in transcription is phase-advanced relative to the RNA-seq data. Because the nighttime decreases are coincident, this makes for a broader BuTT-Seq plateau between ZT10 and ZT14 relative to the RNA-seq data (Figure 7B; green arrows). Remarkably, this comparison is identical to what we reported in a comparison between a nuclear run-on assay of per transcription and a RNase protection assay of steady-state per mRNA (So & Rosbash, 1997). Those assays were low throughput, limiting the number of genes that could be examined. Nonetheless, *tim* was assessed in parallel and was more equivocal, i.e., it did not show such a striking distinction between transcription and mRNA levels (So & Rosbash, 1997); this is similar to what we observe here between tim BuTT-Seq and RNA-seq data (Figure 7B). The parallels between these two different assays done twenty-five years apart indicate that the circadian regulation of *per* is quantitatively and perhaps even qualitatively unusual compared to the other core clock genes and most importantly support the assertion that the BuTT-Seq exon signal successfully assays transcription.

## DISCUSSION

BuTT-Seq is a facile method for the analysis of nascent RNA and sequences the 3’-ends of chromatin associated RNA. In most analyzed reads, the 3’ end is the base most recently synthesized by RNAPII. Although this same strategy is employed by a wide range of methods, BuTT-Seq offers a much simpler workflow: it can produce libraries from purified nascent RNA in as few as six hours and is reproducible down to 10,000 cells.

Although the choice of TGIRT for reverse transcription and adaptor ligation is the most notable difference from these previous methods, many of them employ selection steps that BuTT-seq does not, for example phosphorylation-state specific immunoprecipitation of RNAPII. scRNA-seq and 3NT-seq also deplete uncapped RNA species with a cap-sensitive exonuclease. Although BuTT-Seq circumvents this step, it nevertheless correlates well with 3NT-Seq (Figure 1B and 1C), suggesting that cap-selection may not have a large effect on signal in regions of interest. Relative to these earlier methods, BuTT-seq bears the greatest resemblance to hNET-seq, which also does not incorporate a selection step. Despite the lack of a polymerase-bound or capped RNA selection in the BuTT-Seq protocol, it shows similar signals at protein-coding genes with a much simplified workflow and improved input requirements.

The capacity of BuTT-Seq to address transcriptional pausing detail is highlighted in experiments demonstrating that the transcriptional elongation and SEC inhibitor KL-1 causes transcriptional pause sites to recede towards the TSS. This observation implies the existence of multiple pausing checkpoints downstream from the TSS, the closest of which is rapidly resolved by the recruitment of the SEC shortly after transcriptional initiation. The existence of multiple pausing checkpoints resembles the recently proposed “pausing zone” – a region of enhanced RNA polymerase pausing – adjacent to the transcription start site (Fong et al., 2022). Notably, this paper reported a phosphorylation mutant of Spt5 – Spt5 KOWx4-KOW5 – caused the “pausing zone” to shrink towards the TSS in mammalian cells. This pausing zone finding corroborates our observation of multiple pause sites and suggests that this mode of pause-release regulation may be conserved between insects and mammals. Unlike the KOWx4-KOW5 mutant, however, we found that changes in pausing were primarily dominated by shifting of discrete major pause sites. Whether this difference is driven by a difference in analysis or represents a species-specific phenomenon remains to be seen.

These pause sites have an intriguing relationship to subnucleosomal fragments obtained through MNase-Seq: increased proximity of the pause site to the -1 nucleosome center is negatively correlated with sensitivity to KL-1 treatment and positively correlated with a loss of contact with the distal half of the nucleosomal dyad. These differences in nucleosomal contacts may be a consequence of RNAPII proximity as previously suggested(Ramachandran et al., 2017). However, the insensitivity to KL-1 in genes with pause sites directly adjacent to the center of the nucleosome suggests more complex regulation. Surprisingly, KL-1 appears to also cause transcriptional deceleration near the TSS in genes containing H2Av in the +1 nucleosome, which is depleted of promoter-proximal pausing in *Drosophila*(Weber et al., 2014). This suggests that recruitment of the SEC might occur concurrently with transcriptional initiation at H2Av-marked genes, which then bypasses promoter-proximal pausing.

A strength of BuTT-Seq should be its ability to interrogate samples where input material may be limited, like primary tissue. To this end, we assayed fly heads with BuTT-Seq at 6 different timepoints and focused on 4 clock genes. *per* features a very prominent stable pause signal as previously observed. Future experiments should indicate whether the rhythmic transcription of *per* is regulated at least in part through pause release. In this context, there is evidence that *per* mRNA is more stable early in its accumulation cycle (So et al.). This is when pause release should be prominent, before the large RNAPII increase in the elongating region. Perhaps there is a mRNP difference between transcripts due to pause release compared to those that accumulate later due to increased transcriptional initiation.

In this context, we have previously explored the relative contributions of transcriptional vs. post-transcriptional regulation to circadian mRNA oscillations (e.g., REF). Given that the BuTT-Seq signal along the gene body represents an uncommon approach towards examining transcriptional dynamics – essentially, the quantification of RNAPII density – we revisited this question in a simple way by examining the relationship between BuTT-Seq signal and mRNA-Seq signal. Interestingly, the amplitude, period, and phase of the BuTT-Seq signal and the mRNA-seq signal were remarkably similar – with BuTT-Seq phase leading mRNA-Seq by no more than ∼1 hour. Interestingly, we previously reported a “hump” of *per* transcription that precedes the peak of per mRNA; comparing the two curves suggested interesting *per* post-transcriptional regulation in addition to its more expected transcriptional regulation (So & Rosbash, 1997). The same “hump” is visible in BuTT-Seq now 25 years later (Figure 6B). Although further exploration is beyond the scope of this study, these observations as well as the paused polymerase peak in the *per* promoter region (Figure 6A and Abruzzi et al. 2013(Taylor & Hardin, 2008)) suggest that *per* is subject to one or more unusual modes of regulation relative to other Clk-regulated core circadian genes.

The extent of tissue-specific transcriptional pausing and pause release has received relatively little attention. The relationship between pausing and elongation was non-linear in differentially expressed genes between heads and S2 cells; in other words, differences in pausing signal was frequently not accompanied by a concordant change in gene body signal. This finding echoes a previous study analyzing pausing through RNAPII ChIP-Seq in different mammalian tissues(Day et al., 2016). One possibility is that pausing enables a second layer of regulation, wherein the combinatorial recruitment of initiation and pause release factors enable tissue-specific gene expression(Adelman & Lis, 2012). The discordant relationship we report implies that pause release may be a rate-limiting step in the expression of many genes including *per*, and that specific tissue-specific factors may be required to enable pause-release and the transition from pausing to processive elongation. Such factors are a subject of considerable interest. In any case, we suggest the BuTT-Seq will provide future insights into transcriptional regulation from many other areas of investigation well beyond flies and circadian biology.

## Acknowledgements

We thank Kate Abruzzi for suggestions, Corrie Ratner and Evelyn Keefer for conducting sequencing, and Yuki Aoi, Edwin R. Smith, and Ali Shilatifard for providing KL-1. We thank Kristina Zumer and Patrick Cramer for sharing code used for data visualization.

## Author Contributions

A.D.Y. conceived of the study under the guidance of M.R. A.D.Y performed the experiments and data analysis. A.D.Y. and M.R. wrote the manuscript.

## Methods

### S2 cell culture and KL-1 treatment

S2 cells were obtained from the ATCC (CRL-1963) and cultured in Schneiders Medium supplemented with 10% FetalGro (rmbio, FGX-BBT) and 1% Penicillin/Streptomycin (ThermoFisher, 15140122).

### Nascent RNA isolation in S2 cells

Nascent RNA isolation was adapted from (Khodor et al., 2012). S2 cells grown in a T25 flask were harvested by scraping and pelleted in a centrifuge at 900g for 5 minutes, washing once with 10ml 1x cold PBS. Unless otherwise specified, 5x10^5^ cells were used. Washed cells were resuspended in 500µl cell lysis buffer (10mM Tris pH7.5, 2mM MgCl_2_, 10mM kCl, 0.6mM spermidine, 0.2mM spermine, 3mM TCEP, 0.03% Tween-20, and 0.1% BSA) and transferred into 2ml dounce homogenizers. Cells were homogenized with 10 strokes with Pestle A and 15 strokes with Pestle B. Homogenized cells were passed through a 10µM filter (Sysmex, 04-0042-2314) and centrifuged at 1000g for 5 minutes. Supernatant was removed and nuclei were resuspended in 100µl nuclear lysis buffer (10mM HEPES-KOH pH7.6, 100mM KCl, 0.1mM EDTA, 10% Glycerol, 0.15mM Spermine, 0.5mM Spermidine, 0.1mM NaF, 0.1mM Na_3_VO_4_, 0.1mM ZnCl_2_, 1mM TCEP, 0.1 units/µl SUPERaseIn, and 1x cOmplete protease inhibitor) and placed on a Thermomix set to 4°C and shaking at 1400 RPM. 100µl NUN buffer (25mM HEPES-KOH, pH7.6, 300mM NaCl, 1M Urea, 1% NP-40, 1mM TCEP, 3% Empigen, 0.1 units/µl SUPERaseIn, 1x cOmplete protease inhibitor) was added dropwise while shaking. Tubes were capped and left shaking at 4°C for 10 minutes. Chromatin was pelleted in the centrifuge at 21000g for 10 minutes, and the supernatant was discarded. The resulting chromatin pellet was resuspended in 500µl TRIzol and incubated at 60°C for 10 minutes, then transferred to a phase lock tube. RNA was then isolated with chloroform according to standard procedure and resuspended in a 10µl Turbo DNAse reaction containing 1x Turbo DNase Buffer and 1.5µl Turbo DNase, and DNase treatment was done for 30 minutes according to protocol. 2µl DNase treated RNA was used for BuTT-Seq.

### Nascent RNA isolation in fly heads

Flies were frozen on dry ice. 5 heads were removed on dry ice and transferred into a 2ml dounce homogenizer. 500µl nuclear lysis buffer was added to each homogenizer, and heads were dounced for 10 strokes with Pestle A and 20 strokes with Pestle B. The resulting homogenate was first filtered through a 20µM filter, then a 20µM filter. Nuclei were spun down for 1000g for 5 minutes and washed once with 500µl nuclear lysis buffer. Resulting nuclei were subject to chromatin isolation as with S2 cells.

### BuTT-Seq library prep

BuTT-Seq library prep consists of two consecutive TGIRT protocols for first and second strand synthesis, modified to incorporate an UMI and minimize input.

To prepare SCR2R primer, 10µl 10µM R2 RNA (rArArGrArUrCrGrGrArArGrArGrCrArCrArCrGrUrCrUrGrArArCrUrCrCrArGrUrCrArC/3Sp C3/) was mixed with 10µl 10µM SCR2 DNA (CAAGCAGAAGACGGCATACGAGATNNNNNNNNGTGACTGGAGTTCAGACGTGTGCTCTTCCGATCTTN), where N is a hand-mixed equimolar ratio of A/T/C/G, along with 5µl 10x annealing buffer (10mM Tris-HCl, pH7.5, 10mM EDTA, 10mM TCEP pH 7.5) and 25µl H_2_O. SCR2R primer was ordered as PAGE-purified, while SCR2 DNA was ordered as a PAGE-purified Ultramer from IDT. Primers were added to a pre-heated 88°C thermal cycler, incubated for 2 minutes, and then cooled to 10°C at a rate of 0.1°C/second and held at 10°C.

To prepare MER1R primer, 10µl 10µM R1ME RNA (rCrUrGrUrCrUrCrUrUrArUrArCrArCrArUrCrUrGrArCrGrCrUrGrC/3SpC3/) was mixed with 10µl 10µM R1ME DNA (GCAGCGTCAGATGTGTATAAGAGACAGN), where N is a hand-mixed equimolar ratio of A/T/C/G, along with 5µl 10x annealing buffer (10mM Tris-HCl, pH7.5, 10mM EDTA, 10mM TCEP pH 7.5) and 25µl H_2_O. Both primers were PAGE-purified from IDT. Primers were added to a pre-heated 88°C thermal cycler, incubated for 2 minutes, and then cooled to 10°C at a rate of 0.1°C/second and held at 10°C.

All primers were aliquoted into single-used aliquots and stored at -80°C for up to 6 months. Reaction master mixes were prepared with 0.5µl 10x TGIRT Buffer(100mM HEPES pH8, 500mM NaCl, 50mM MgCl2, 10mM TCEP), 0.2µl SCR2R, 0.2µl TGIRT, and 1.1µl 50% PEG3350. Master mixes were prepared for at least 4 reactions at a time to minimize pipetting errors. To ensure the reactions were well-mixed, reactions were stirred ten times with a pipette tip, and flicked and spun down twice. 2µl of master mix was added to 2µl RNA in a PCR strip tube, flicking and spinning down twice to mix. Assembled reactions were incubated on ice for 30 minutes.

While incubating, 1µl 5mM dNTPs was mixed with 30µl 2M NaCl. After 30 minutes, 1µl dNTPs/NaCl mixture was added to TGIRT reactions and placed on the thermal cycler with the following program: 25°C for 1 minute, ramping to 60°C at 1°C/second, 60°C for 10 seconds, then holding at 4°C. Reactions were moved on to ice and 2µl Exonuclease III, 2µl H_2_O, and 1µl NEBuffer 1 were added and flicked to mix. Reactions were incubated at 37°C for 8 minutes, then moved onto ice. Exonuclease III treatment partially digests unused primers. This step is unnecessary with sufficient input, but with low input samples, excess primers may be overamplified and contaminate the final library.

To hydrolyze the RNA, 3µl 1.3M NaOH was added to each sample and heated at 95°C for 5 minutes. After cooling to room temperature, 3µl 1.3M HCl was added to each sample to neutralize the reaction. 24µl 95% EtOH and 24µl Ampure XP beads were added to each sample, flicked to mix, and left to incubate for 10 minutes at room temperature. Beads were washed twice with 200µl 80% EtOH and resuspended in 4.2µl H_2_O and 4µl supernatant was moved to fresh PCR tubes.

Second-strand synthesis master mixes were prepared with 1µl TGIRT Buffer, 0.3µl MER1R, 0.3µl TGIRT, 2µl 50% PEG3350, and 0.4µl H_2_O. Master mixes were mixed as described earlier. 4µl master mix was added to cDNA samples, mixed as described earlier, and assembled reactions were left to incubate on ice for 30 minutes.

While incubating, 1µl 20mM dNTPs was mixed with 10µl 2M NaCl. After 30 minutes, 2µl dNTPs/NaCl mixture was added to TGIRT and placed on the thermal cycler with the following program: 25°C for 1 minute, ramping to 60°C at 1°C/second, 60°C for 30 seconds, then holding at 4°C. Reactions were moved on to ice, and 1µl 0.2% SDS was added to each reaction.

Reactions were incubated 55°C, then moved to room temperature. 30µl Ampure XP beads were added to each reaction, incubated for 5 minutes at room temperature, washed twice, and then eluted in 9µl H_2_O.

dsDNA was added to PCR reactions containing 25µl 2x NEBNext Ultra II Q5 Master mix, 2.5µl P7 primer (CAAGCAGAAGACGGCATACGAG), 2.5µl barcoded Ad1 primer(AATGATACGGCGACCACCGAGATCTACACTCGTCGGCAGCGTCAG ATGTGTAT), and 11µl H_2_O. The assembled PCR reaction was subject to the following PCR program: 72°C for 3 minutes, 98°C for 30 seconds, 9-15 cycles of 98°C for 15 seconds and 65°C for 30 seconds, a final extension of 72°C for 2 minutes, and hold at 10°C.

PCR reactions were cleaned with 65µl Ampure XP beads following the standard protocol and resuspended in 10µl 1x Purple Loading Dye. Samples were loaded onto an 8% TBE gel alongside 1µl of TriDye Ultra Low Range DNA ladder mixed with 9µl 1x Purple Loading Dye and run at 180v for 45 minutes. Gels were post-stained with 1x Sybr Gold for 5 minutes, and the smear above 150nt were excised into a 0.5ml tube. The maximum fragment size produced in this protocol should be around ∼700nt.

0.5ml tubes had a hole poked in the bottom with a 21-gauge needle and nested inside a 2.0ml tube, and centrifuged for 21000g for 3 minutes. If any gel fragments remained in the 0.5ml tube, a second hole was poked and centrifuged again. 600µl 300mM NaCl was added to each tube, and tubes were incubated at 75°C for 15 minutes or 4°C overnight, preferably with agitation or shaking. Supernatant and gel fragments were transferred into a Costar Spin-X Centrifuge Tube Filter with a wide-bore 1000µl pipette tip, or a 1000µl pipette tip with the tip cut off, and spun for 2 minutes at 21000g. Filters were discarded and 600µl isopropanol and 0.7µl GlycoBlue was added to each tube, and left to incubate for at least 15 minutes at room temperature or in -20°C for up to overnight. Samples were spun at 21000g for 30 minutes, washed twice with 80% EtOH, and resuspended in 7-10µl MilliQ H_2_O. 2µl was used for analysis on a Tapestation 4200 D1000 tape and quantified using peak quantification centered around 170nt. Samples were sequenced on a NextSeq 500 with at least 8bp dual index reads.

### BuTT-Seq Data Processing

A Snakemake file for analyzing BuTT-Seq is available at, which also includes all custom scripts. Prior to demultiplexing, RunInfo.xml was altered so that Read 2 is not counted as an Indexed read, and samples were demultiplexed using the i5 index. This will output three files: Read 1 corresponds to Read 1, Read 2 corresponds to the UMI, and Read 3 corresponds to Read UMIs were extracted from Read 2 and appended to Read 1 and 3 individually using umi_tools extract with the following command: extract -I READ2_UMI --extract-method=regex --bc-pattern=’^(?P.<umi_1>{{8}})’ --read2-in=READ1_OR_3 -- stdout=umiextracted_reads/filler.fq.gz --read2-out=OUTPUT.fq.gz (Smith et al., 2017). This will also produce a filler file, which can be deleted. Reads were trimmed using fastp and the following settings: -trim_poly_g --trim_poly_x -F 1 --adapter_sequence AAGATCGGAAGAGC --adapter_sequence_r2 CTGTCTCTTATA (Chen et al., 2018). Reads were subject to 2-pass mapping with STAR to dm6 using the following settings: --alignMatesGapMax 100000 -- outSAMstrandField intronMotif --outFilterMismatchNoverLmax 0.05 --outFilterMultimapNmax 1 --outSJfilterReads Unique. Aligned reads were converted to BAM files with Samtools, and deduplicated using umi_tools dedup (Li et al., 2009). Small RNAs and exon ends were removed using SAM tools and a custom bed file, and soft clipping was removed using a custom script (Breese & Liu, 2013).

### Pro-Seq and 3NT-Seq Data Processing

Fastq files were obtained from the SRA using fasterq-dump and aligned to dm6 using STAR with the following settings: --alignMatesGapMax 50000 --outFilterMismatchNoverLmax 0.06 -- outFilterMatchNmin 15 --outFilterMultimapNmax 1 --outSJfilterReads Unique. Aligned reads were converted to BAM files with Samtools (Li et al., 2009). Reads were truncated to the first nucleotide of Read 2 using a custom script.

### ChIP-Seq Data Processing and Peak Calling

Fastq files were retrieved from the SRA using fasterq-dump and aligned to dm6 with bwa mem. Peaks were called using MACS2 using the following parameters: --nomodel --extsize 200 -q 0.01.

### BuTT-Seq Pause Analysis

Reads were truncated to the first nucleotide of Read 2 using a custom script and converted into stranded bedgraph files using Deeptools (Nojima et al., 2016; Ramirez et al., 2014). Bedgraph files were restricted to the first 200nt downstream of each TSS genome-wide, with overlapping genes being merged together. Single-nucleotide peaks were called using the Pause-Detection Algorithm (PDA) (Gajos et al., 2021). Single-nucleotide peaks were used as a reference for quantifying single-nucleotide BAM files using FeatureCounts, and multiple peaks in a single gene were filtered for the highest peak in R (Liao et al., 2014).

Rather than calculating pausing indices using fixed regions, pausing regions were calculated with reference to highest pause peak in each gene. Pause regions were defined as the region from the TSS to the highest pause, and elongation regions were defined as 1000bp downstream of the highest pause.

### mNase-Seq analysis

Fastq files were retrieved from the SRA using fasterq-dump and aligned to dm6 with bwa mem using default settings. BAM files were filtered by insert size using SAMtools. Nucleosome positions were called as previously described, except rather than filtering for the downstream nucleosome, entire alignments were first used for unbiased analysis. Following unbiased analysis, the -1 nucleosome was described as any nucleosome that overlapped an annotated TSS. Bedtools was used to determine nucleosome centers and to calculate the distance to the nearest BuTT-Seq pause peak (Quinlan, 2014).

### H2Av ChIP-Seq analysis

Peaks were called using MACS2 using the following parameters: --nomodel --extsize 200 -q 0.01 (Zhang et al., 2008).

### RNA-Seq analysis

Fastq files were retrieved from the SRA using fasterq-dump and aligned to dm6 using STAR with the following settings: --alignMatesGapMax 50000 --outFilterMismatchNoverLmax 0.06 -- outFilterMatchNmin 15 --outFilterMultimapNmax 1 --outSJfilterReads Unique. Aligned reads were converted to BAM files with Samtools (Li et al., 2009). Reads were counted using featurecounts using Refseq genes as a reference (Liao et al., 2014). Normalized count files were generated using DESeq2 (Love et al., 2014). For each gene, reads across all timepoints were normalized to the highest timepoint and double-plotted using ggplot2 (Wickham, 2016).

### Metagene profiles and gene plots

Mapped reads in each sequencing experiment were used to generate normalization factors with DESeq2 using counts at single nucleotide peaks as a reference (Love et al., 2014). Normalization factors were used to convert mapped reads into coverage plots using deepTools (Ramirez et al., 2014). To simplify visualization, only plus strand reads and genes were used. Coverage plots were scaled to specified regions using deepTools computematrix with a bin size of 1, and the output was used for further processing in R.

In R, outliers defined as the top and bottom 0.1% regions were removed, a pseudo-count of 1 was added to all positions and converted into log_2_, from which means and 95% confidence intervals were determined using the bootstrap method with 10,000 repetitions. Means and confidence intervals were plotted using ggplot2 ∂(Wickham, 2016).

For gene-specific plots and some figure building, Plotgardener was used (Kramer et al., 2022).

## REFERENCES

Abruzzi, K. C., Rodriguez, J., Menet, J. S., Desrochers, J., Zadina, A., Luo, W., Tkachev, S., & Rosbash, M. (2011). Drosophila CLOCK target gene characterization: implications for circadian tissue-specific gene expression. Genes Dev, 25(22), 2374–2386. https://doi.org/10.1101/gad.174110.11110.1101/gad.178079.111

Adelman, K., & Lis, J. T. (2012). Promoter-proximal pausing of RNA polymerase II: emerging roles in metazoans. Nat Rev Genet, 13(10), 720–731. https://doi.org/10.1038/nrg3293

Breese, M. R., & Liu, Y. (2013). NGSUtils: a software suite for analyzing and manipulating next-generation sequencing datasets. Bioinformatics, 29(4), 494–496. https://doi.org/10.1093/bioinformatics/bts731

Chen, S., Zhou, Y., Chen, Y., & Gu, J. (2018). fastp: an ultra-fast all-in-one FASTQ preprocessor. Bioinformatics, 34(17), i884–i890. https://doi.org/10.1093/bioinformatics/bty560

Churchman, L. S., & Weissman, J. S. (2012). Native elongating transcript sequencing (NET-seq). Curr Protoc Mol Biol, Chapter 4, Unit 4 14 11-17. https://doi.org/10.1002/0471142727.mb0414s98

Core, L. J., Waterfall, J. J., & Lis, J. T. (2008). Nascent RNA sequencing reveals widespread pausing and divergent initiation at human promoters. Science, 322(5909), 1845–1848. https://doi.org/10.1126/science.1162228

Day, D. S., Zhang, B., Stevens, S. M., Ferrari, F., Larschan, E. N., Park, P. J., & Pu, W. T. (2016). Comprehensive analysis of promoter-proximal RNA polymerase II pausing across mammalian cell types. Genome Biol, 17(1), 120. https://doi.org/10.1186/s13059-016-0984-2

Fong, N., Sheridan, R. M., Ramachandran, S., & Bentley, D. L. (2022). The pausing zone and control of RNA polymerase II elongation by Spt5: Implications for the pause-release model. Mol Cell, 82(19), 3632–3645 e3634. https://doi.org/10.1016/j.molcel.2022.09.001

Gajos, M., Jasnovidova, O., van Bommel, A., Freier, S., Vingron, M., & Mayer, A. (2021). Conserved DNA sequence features underlie pervasive RNA polymerase pausing. Nucleic Acids Res, 49(8), 4402–4420. https://doi.org/10.1093/nar/gkab208

He, Q., Johnston, J., & Zeitlinger, J. (2015). ChIP-nexus enables improved detection of in vivo transcription factor binding footprints. Nat Biotechnol, 33(4), 395–401. https://doi.org/10.1038/nbt.3121

Judd, J., Wojenski, L. A., Wainman, L. M., Tippens, N. D., Rice, E. J., Dziubek, A., Villafano, G. J., Wissink, E. M., Versluis, P., Bagepalli, L., Shah, S. R., Mahat, D. B., Tome, J. M., Danko, C. G., Lis, J. T., & Core, L. J. (2020). A rapid, sensitive, scalable method for Precision Run-On sequencing (PRO-seq). bioRxiv, 2020.2005.2018.102277. https://doi.org/10.1101/2020.05.18.102277

Khodor, Y. L., Menet, J. S., Tolan, M., & Rosbash, M. (2012). Cotranscriptional splicing efficiency differs dramatically between Drosophila and mouse. RNA, 18(12), 2174–2186. https://doi.org/10.1261/rna.034090.112

Kramer, N. E., Davis, E. S., Wenger, C. D., Deoudes, E. M., Parker, S. M., Love, M. I., & Phanstiel, D. H. (2022). Plotgardener: cultivating precise multi-panel figures in R. Bioinformatics, 38(7), 2042–2045. https://doi.org/10.1093/bioinformatics/btac057

Kuintzle, R. C., Chow, E. S., Westby, T. N., Gvakharia, B. O., Giebultowicz, J. M., & Hendrix, D. A. (2017). Circadian deep sequencing reveals stress-response genes that adopt robust rhythmic expression during aging. Nat Commun, 8, 14529. https://doi.org/10.1038/ncomms14529

Li, H., Handsaker, B., Wysoker, A., Fennell, T., Ruan, J., Homer, N., Marth, G., Abecasis, G., Durbin, R., & Genome Project Data Processing, S. (2009). The Sequence Alignment/Map format and SAMtools. Bioinformatics, 25(16), 2078–2079. https://doi.org/10.1093/bioinformatics/btp352

Liang, K., Smith, E. R., Aoi, Y., Stoltz, K. L., Katagi, H., Woodfin, A. R., Rendleman, E. J., Marshall, S. A., Murray, D. C., Wang, L., Ozark, P. A., Mishra, R. K., Hashizume, R., Schiltz, G. E., & Shilatifard, A. (2018). Targeting Processive Transcription Elongation via SEC Disruption for MYC-Induced Cancer Therapy. Cell, 175(3), 766–779e717.https://doi.org/10.1016/j.cell.2018.09.027

Liao, Y., Smyth, G. K., & Shi, W. (2014). featureCounts: an efficient general purpose program for assigning sequence reads to genomic features. Bioinformatics, 30(7), 923–930. https://doi.org/10.1093/bioinformatics/btt656

Love, M. I., Huber, W., & Anders, S. (2014). Moderated estimation of fold change and dispersion for RNA-seq data with DESeq2. Genome Biol, 15(12), 550. https://doi.org/10.1186/s13059-014-0550-8

Mahat, D. B., Kwak, H., Booth, G. T., Jonkers, I. H., Danko, C. G., Patel, R. K., Waters, C. T., Munson, K., Core, L. J., & Lis, J. T. (2016). Base-pair-resolution genome-wide mapping of active RNA polymerases using precision nuclear run-on (PRO-seq). Nat Protoc, 11(8), 1455–1476. https://doi.org/10.1038/nprot.2016.086

Mayer, A., di Iulio, J., Maleri, S., Eser, U., Vierstra, J., Reynolds, A., Sandstrom, R., Stamatoyannopoulos, J. A., & Churchman, L. S. (2015). Native elongating transcript sequencing reveals human transcriptional activity at nucleotide resolution. Cell, 161(3), 541–554. https://doi.org/10.1016/j.cell.2015.03.010

Mayer, A., Landry, H. M., & Churchman, L. S. (2017). Pause & go: from the discovery of RNA polymerase pausing to its functional implications. Curr Opin Cell Biol, 46, 72–80. https://doi.org/10.1016/j.ceb.2017.03.002

Mohr, S., Ghanem, E., Smith, W., Sheeter, D., Qin, Y., King, O., Polioudakis, D., Iyer, V. R., Hunicke-Smith, S., Swamy, S., Kuersten, S., & Lambowitz, A. M. (2013). Thermostable group II intron reverse transcriptase fusion proteins and their use in cDNA synthesis and next-generation RNA sequencing. RNA, 19(7), 958–970. https://doi.org/10.1261/rna.039743.113

Nechaev, S., Fargo, D. C., dos Santos, G., Liu, L., Gao, Y., & Adelman, K. (2010). Global analysis of short RNAs reveals widespread promoter-proximal stalling and arrest of Pol II in Drosophila. Science, 327(5963), 335–338. https://doi.org/10.1126/science.1181421

Nojima, T., Gomes, T., Carmo-Fonseca, M., & Proudfoot, N. J. (2016). Mammalian NET-seq analysis defines nascent RNA profiles and associated RNA processing genome-wide. Nat Protoc, 11(3), 41©3-428. https://doi.org/10.1038/nprot.2016.012

Nojima, T., Gomes, T., Grosso, A. R. F., Kimura, H., Dye, M. J., Dhir, S., Carmo-Fonseca, M., & Proudfoot, N. J. (2015). Mammalian NET-Seq Reveals Genome-wide Nascent Transcription Coupled to RNA Processing. Cell, 161(3), 526–540. https://doi.org/10.1016/j.cell.2015.03.027

Quinlan, A. R. (2014). BEDTools: The Swiss-Army Tool for Genome Feature Analysis. Curr Protoc Bioinformatics, 47, 11 12 11–34. https://doi.org/10.1002/0471250953.bi1112s47

Ramachandran, S., Ahmad, K., & Henikoff, S. (2017). Transcription and Remodeling Produce Asymmetrically Unwrapped Nucleosomal Intermediates. Mol Cell, 68(6), 1038–1053 e1034. https://doi.org/10.1016/j.molcel.2017.11.015

Ramirez, F., Dundar, F., Diehl, S., Gruning, B. A., & Manke, T. (2014). deepTools: a flexible platform for exploring deep-sequencing data. Nucleic Acids Res, 42(Web Server issue), W187-191. https://doi.org/10.1093/nar/gku365

Rodriguez, J., Tang, C. H., Khodor, Y. L., Vodala, S., Menet, J. S., & Rosbash, M. (2013). Nascent-Seq analysis of Drosophila cycling gene expression. Proc Natl Acad Sci U S A, 110(4), E275–284. https://doi.org/10.1073/pnas.1219969110

Schlackow, M., Nojima, T., Gomes, T., Dhir, A., Carmo-Fonseca, M., & Proudfoot, N. J. (2017). Distinctive Patterns of Transcription and RNA Processing for Human lincRNAs. Mol Cell, 65(1), 25–38. https://doi.org/10.1016/j.molcel.2016.11.029

Smith, T., Heger, A., & Sudbery, I. (2017). UMI-tools: modeling sequencing errors in Unique Molecular Identifiers to improve quantification accuracy. Genome Res, 27(3), 491–499. https://doi.org/10.1101/gr.209601.116

So, W. V., & Rosbash, M. (1997). Post-transcriptional regulation contributes to Drosophila clock gene mRNA cycling. EMBO J, 16(23), 7146–7155. https://doi.org/10.1093/emboj/16.23.7146

Sousa-Luis, R., Dujardin, G., Zukher, I., Kimura, H., Weldon, C., Carmo-Fonseca, M., Proudfoot, N. J., & Nojima, T. (2021). POINT technology illuminates the processing of polymerase-associated intact nascent transcripts. Mol Cell, 81(9), 1935–1950 e1936. https://doi.org/10.1016/j.molcel.2021.02.034

Taylor, P., & Hardin, P. E. (2008). Rhythmic E-box binding by CLK-CYC controls daily cycles in per and tim transcription and chromatin modifications. Mol Cell Biol, 28(14), 4642–4652. https://doi.org/10.1128/MCB.01612-07

Weber, C. M., Ramachandran, S., & Henikoff, S. (2014). Nucleosomes are context-specific, H2A.Z-modulated barriers to RNA polymerase. Mol Cell, 53(5), 819–830. https://doi.org/10.1016/j.molcel.2014.02.014

Wickham, H. (2016). Ggplot2 : elegant graphics for data analysis. Springer Science+Business Media, LLC.

Wissink, E. M., Vihervaara, A., Tippens, N. D., & Lis, J. T. (2019). Nascent RNA analyses: tracking transcription and its regulation. Nat Rev Genet, 20(12), 705–723. https://doi.org/10.1038/s41576-019-0159-6

Wuarin, J., & Schibler, U. (1994). Physical isolation of nascent RNA chains transcribed by RNA polymerase II: evidence for cotranscriptional splicing. Mol Cell Biol, 14(11), 7219–7225. https://doi.org/10.1128/mcb.14.11.7219-7225.1994

Zhang, Y., Liu, T., Meyer, C. A., Eeckhoute, J., Johnson, D. S., Bernstein, B. E., Nusbaum, C., Myers, R. M., Brown, M., Li, W., & Liu, X. S. (2008). Model-based analysis of ChIP-Seq (MACS). Genome Biol, 9(9), R137. https://doi.org/10.1186/gb-2008-9-9-r137

